# A role for SHARPIN in platelet linear protein ubiquitination and function

**DOI:** 10.1101/2021.01.13.426403

**Authors:** S. F. Moore, X. Zhao, S. Mallah, A.W. Poole, S. J. Mundell, J. L. Hutchinson, I. Hers

## Abstract

SHARPIN (Src homology 3 and multiple ankyrin repeat domains protein (SHANK)- associated RH domain-interacting protein) as part of the linear ubiquitin chain assembly complex (LUBAC) catalyses the addition of linear (Met1-linked) ubiquitin chains to substrates. As part of this complex SHARPIN acts as a multi-functional modulator of immune/inflammatory responses through regulation of NfkB activation. In addition, SHARPIN can act as a negative regulator of integrin function. Despite platelets being anucleate cells several studies have determined potential roles for both ubiquitination and NfkB in regulating platelet function. However, little is known about either linear ubiquitination and/or SHARPIN in mouse platelets. In this study, we evaluated platelet function in mice with impaired SHARPIN expression. We confirmed that SHARPIN was expressed in platelets from wild-type mice but not in mice homozygous for SHARPIN^cpdm^ allele (cpdm/cpdm) and that this correlated with a reduction in linear ubiquitination. Platelet function in response to thrombin was unaffected. In contrast, CRP-XL-and U46619-mediated platelet responses and thrombus formation under flow on a collagen-coated surface were significantly reduced in the cpdm/cpdm mice. This was associated with impaired U46619-mediated intracellular signalling as well as a reduction in CRP-mediated ERK phosphorylation. Despite the reported role for NfκB in regulating platelet function, inhibiting IκBα phosphorylation did not recapitulate the cpdm/cpdm phenotype. Together, these data indicate that the lack of SHARPIN and linear ubiquitination results in impaired thrombosis and platelet functional responses to CRP and U46619. This phenotype is independent of NfκB pathway inhibition but may involve alternative signalling pathways regulated by linear ubiquitination.

**Key Points:**

- SHARPIN plays an essential role in platelet linear protein ubiquitination and CRP and U46619-mediated platelet function
- In vitro thrombosis is significantly impaired in SHARPIN deficient mice

## INTRODUCTION

SHARPIN (Src homology 3 and multiple ankyrin repeat domains protein (SHANK)- associated RH domain-interacting protein) is a 387-amino acid protein originally identified as a SHANK-binding protein enriched in the postsynaptic density (PSD) of excitatory synapses in brain^1^. It is more widely known for its role as a novel multi-functional modulator of immune and inflammatory responses, with a spontaneous null mutation in the SHARPIN gene resulting in a complex inflammatory phenotype in mice. This phenotype is characterised by chronic proliferative dermatitis (cpdm), multi-organ inflammation, eosinophilic infiltration, dysregulated secondary lymphoid organ development and increased expression of type 2 cytokines^2,3^, with notably similarities to human hypereosinophilic syndromes.

The molecular mechanism whereby loss of SHARPIN results in this inflammatory phenotype has been proposed to be due to its role in regulating the canonical activation of NfkB^4,5^ as part of the linear ubiquitin chain assembly complex (LUBAC). This complex consists of SHARPIN, HOIL-1L (RBCK1) and the ubiquitin ligase HOIP (RNF31). LUBAC can regulate a diverse range of cell signalling pathways by catalysing the addition of linear (Met1-linked) ubiquitin chains to substrates. Substrates known to be targets for linear ubiquitination include NEMO (IKKγ/IKK3), RIP1 and RIP2. Linear ubiquitination of NEMO induces multimerization of the IκB kinase complex (IKK) resulting in the phosphorylation and activation of IKKβ (IKK2). This kinase once active phosphorylates NF-kappa-B inhibitor alpha (IκBα) promoting its ubiquitination and subsequent degradation. This degradation of IκBα is required to unmask the nuclear localisation signal of NfkB, freeing NfkB to translocate to the nucleus and activate transcription.

Despite platelets being anucleate cells several studies have determined potential roles for both ubiquitination and NfkB in regulating platelet function. Many proteins are modified by ubiquitination in platelets under both resting and stimulated conditions ^6-9^ and pre-treatment of platelets with inhibitors of either deubiquitinating enzymes or the proteasome reportedly inhibits platelet activation^7,9,10^. Regarding the NfkB pathway, platelets express the functional components required for NfkB activation including the Iκ-kinases and regulatory inhibitor IκBα ^11^. Furthermore, inhibition of IκBα phosphorylation suppresses platelet activation^11,12^. In addition to its role in linear ubiquitination and activation of the NfκB pathway, work in cpdm/cpdm mice has shown that SHARPIN acts as a negative regulator of integrin function by binding to the α-subunit of β_1_ integrins using^13^. Recent work by the group of Shattil using iPS cell-derived platelet-like fragments suggested that SHARPIN can also regulate platelet function by activating the NfκB pathway as well as acting as a negative regulator of integrin α_IIb_β_3_^14^.

In this study, we evaluated the role of SHARPIN and downstream linear ubiquitination in regulating platelet function using cpdm/cpdm mice. Our data demonstrate that loss of functional SHARPIN results in reduced linear ubiquitination and suppression of platelet function. However, this suppression of platelet function is not due to inhibition of the NfkB pathway as inhibition of IκBα phosphorylation failed to recapitulate the phenotype.

## MATERIALS & METHODS

### Materials

FITC-conjugated glycoprotein VI (GPVI, JAQ1, #M011-1), PE-conjugated integrin αIIbβ3 (JON/A, #M023-2), FITC-conjugated CD62P/P-selectin (Wug.E9, #M130-1) and FITC-conjugated integrin α2 (CD49b, GPIa, Sam.C1, #M071-0) anti-mouse antibodies were from Emfret Analytics (Würzburg, Germany). FITC-conjugated integrin β1 (CD29, HM beta 1-1, #MCA2298) and FITC-conjugated integrin β3 (CD61, #MCA2299) were from Bio-Rad (Oxford, UK). CHRONO-LUME® was from Chrono-log Corporation (Labmedics, Manchester, UK). Cross-linked collagen-related peptide (CRP-XL) was synthesized by Prof. Richard Farndale (Department of Biochemistry, University of Cambridge, UK). Bovine fibrinogen was from Enzyme Research Laboratories (South Bend, IN). Collagen Reagens HORM® Suspension was from Takeda (Linz, Austria). FURA-PE3 and Anti-Linear Ubiquitin Antibody, clone LUB9 (#MABS451) were from Millipore (U.K.) Ltd (Watford, UK). Akt (#927), Akt^T308^ (#13038), Akt^S473^ (#4060), ERK1/2^T202/Y204^ (#4370), ERK1/2 (#9102), PLCγ2^Y759^ (#3874), PLCγ2 (#3872), phospho-Ser PKC substrate (#2261), IkBα^S32/36^ (#9246), IkBα (#9242) and IKKα/β^S176/177^ (#2078) antibodies were from Cell Signalling Technologies (London, UK). BMS345541, BI605906, BAY 11-7082 were from Cambridge Bioscience (Cambridge, UK). Takinib was from Selleckchem (Stratech, Newmarket, UK). Enhanced chemiluminescent detection reagents were from GE Healthcare (Bucks, UK). Peroxidase-conjugated and far-red/infrared secondary antibodies were from Jackson ImmunoResearch Ltd (Stratech, Newmarket, UK). NuPAGE SDS–PAGE sample buffer and ActinGreen™ 488 were from Invitrogen (Paisley, UK). cOmplete™ protease inhibitor cocktail tablets were from Roche (West Sussex, UK). All other reagents were from Sigma (Poole, UK), unless otherwise indicated.

### Mice

Animal studies were approved by local research ethics and mice bred for this purpose under a

U.K. Home Office project license (PPL30/3445). C57BL/KaLawRij-SHARPIN^cpdm^/RijSunJ (abbreviated to cpdm) were from The Jackson Laboratory (ME USA). Experiments were performed on age- and sex-matched mice. Mice homozygous for the spontaneous mutation (cpdm/cpdm) have a progressive inflammatory phenotype beginning at 3-5 weeks of age, therefore all experiments using cpdm/cpdm were performed on mice no older than 7 weeks.

### Platelet Isolation

Mice (6–7 weeks old) were sacrificed by rising CO_2_ inhalation, in accordance with Schedule 1 of the Animals (Scientific Procedures) Act (1986). Blood was drawn by cardiac puncture into a syringe containing 4% trisodium citrate (1:9, v/v). Complete blood counts were conducted (Pentra ES60, Horiba) and adjusted for anticoagulant volume. Human blood was obtained with approval from the local Research Ethics Committee of the University of Bristol from healthy drug-free volunteers, who gave full informed consent in accordance with the Declaration of Helsinki. Washed platelets were obtained as previously described ^15^. Platelets were pelleted in the presence of 140 nM PGE1 and 0.02 U/ml apyrase and resuspended at 4 × 10^8^/ml in modified HEPES-Tyrode’s (145 mM NaCl, 3 mM KCl, 0.5 mM Na_2_HPO_4_, 1 mM MgSO_4_, 10 mM HEPES, pH 7.2, 0.1% (w/v) D-glucose, 0.02 U/ml apyrase).

### Flow Cytometry

Flow cytometry analyses were performed as previously described^16^. Integrin α_IIb_β_3_ activation and α-granule secretion were measured using a PE-conjugated antibody (JON/A) directed against the high affinity form of integrin α_IIb_β_3_ and a FITC-conjugated antibody (Wug.E9) for the α-granule marker CD62P (P-selectin). Surface expression of GPVI, integrin α_2_, β_1_, α_IIb_ and β_3_ were measured in resting platelets using FITC-conjugated antibodies. Samples were fixed with 1% paraformaldehyde for 30 min and 10,000 platelet events per sample recorded on a BD Accuri™ C6 Plus.

### δ-Granule Secretion

Agonist-stimulated δ-granule secretion was assessed by monitoring ATP release from platelets (2 × 10^8^/ml) using a luciferin/luciferase reagent (CHRONO-LUME®), calibrated by addition of a known amount of ATP standard at fixed concentration following ligand addition and monitoring. Luminescence was measured using a Tecan Infinite® 200 PRO plate reader as previous described ^17^.

### In Vitro Thrombus Formation

Mice were sacrificed by rising CO_2_ inhalation, in accordance with Schedule 1 of the Animals (Scientific Procedures) Act (1986). Blood was drawn by cardiac puncture into a syringe containing 40 μM PPACK and 5 U/ml clinical-grade heparin. Thrombus formation was performed as previously described^18^ by perfusing DiOC_6_-labelled blood (120 dynes/cm^2^, 3 min) over a Cellix Vena8 biochip (V8GCS-NA-NT-800-80-P10) coated with 50 μg/ml fibrillar HORM® collagen. Biochips were washed and fixed before images captured using a 40× oil immersion objective on a Leica SPE single channel confocal laser scanning microscope attached to a Leica DMi8 inverted epifluorescence microscope. Three randomly chosen fields of view were analysed per sample. Quantification was performed using Volocity® 6.1.1 Quantitation software (Perkin Elmer Inc., San Jose, CA, USA) and ImageJ 1.52k used to capture the representative images.

### Cytosolic [Ca2+] measurements

Changes in cytosolic [Ca^2+^]_i_ were measured in washed platelets loaded with FURA-PE3 using a Tecan Infinite® 200 PRO plate reader. Platelets (4×10^8^/ml) were incubated in CGS buffer (120 mM sodium chloride, 25.8 mM trisodium citrate, 0.1 % (w/v) D-glucose) containing 0.35 % BSA, 0.1 % Pluronic F127 and 4 μM FURA-PE3 (45 min) before pelleting and resuspending in modified HEPES-Tyrode’s containing 1 mM CaCl_2_. Platelets (1×10^8^/ml) were added to a 96 well plate for experiments. Baseline recordings were taken before agonist addition.

### Protein Extraction and Immunoblotting

Platelets (4 × 10^8^/ml) were incubated with agonists as indicated. Platelet suspensions were lysed directly in 4× NuPAGE sample buffer containing 0.5 M DTT. Lysates were analysed by SDS-PAGE/western blotting as previously described ^19^ and visualised by either enhanced chemiluminescence or near-infrared detection (Odyssey®CLx, LI-COR). For quantification of immunoblotting; densitometry was performed using LI-COR Image Studio™ 5 Software.

### Data Analysis

Data were analysed using GraphPad Prism 7 software. All data are presented as the mean ± s.e.mean of at least three independent observations. Data presented with statistical analysis were tested as described in the figure legends.

## RESULTS

### Platelets from cpdm/cpdm mice do not express SHARPIN and have reduced linear ubiquitination

To assess the role of SHARPIN in regulating platelet function we used the C57BL/KaLawRij-SHARPIN^cpdm^/RijSunJ (cpdm) mouse line. The cpdm mutation results in the loss of SHARPIN protein expression due to a single base pair deletion in the 3’ end of exon 1 of SHARPIN, creating a shift in the open reading frame predicted to cause a premature stop codon at position 624 ^2^. In line with this we detected SHARPIN in wild-type platelets (+/+, matched littermates) but not in platelets from mice homozygous for SHARPIN^cpdm^ allele (cpdm/cpdm) (**Fig. 1A**). As part of the LUBAC complex, SHARPIN is involved in catalysing the addition of linear (Met1-linked) ubiquitin chains to substrates with previous studies demonstrating that cpdm/cpdm mice have reduced linear ubiquitination^20^. Using a monoclonal antibody (LUB9), which specifically recognizes linear (Met-1) ubiquitin chains, we observed in wild-type (+/+) platelets basal levels of linear ubiquitination (**Fig. 1B**) which increased upon stimulation. Consistent with the role of SHARPIN in the LUBAC complex, linear ubiquitination in cpdm/cpdm platelets was reduced under both basal and stimulated conditions (**Fig. 1B**).

**Figure 1.**
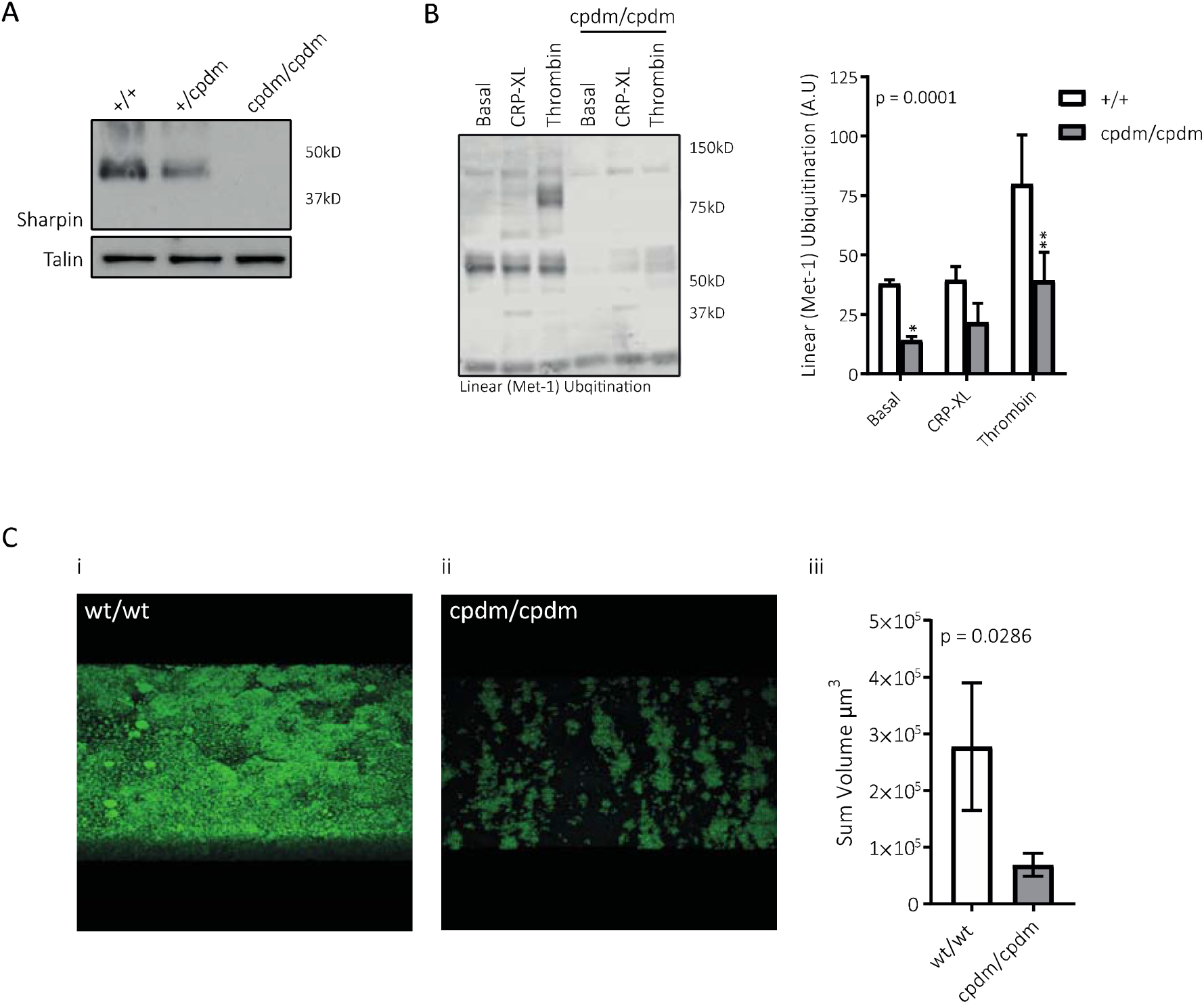
Platelets from cpdm/cpdm mice do not express SHARPIN, have reduced linear ubiquitination and show reduced accumulation on collagen under shear. Washed platelets were treated as indicated before lysis in 4× NuPAGE sample buffer containing 0.5 M DTT before SDS-PAGE/Western blotting. **A)** SHARPIN and TALIN (control) expression in wild-type and cpdm/cpdm platelets. **B)** Detection of linear (Met1-linked) ubiquitination by LUB9 antibody in basal and stimulated wild-type and cpdm/cpdm platelets. **i** representative blot, **ii** summary densitometry n=3 mean +/-s.e., **C)** Whole blood anticoagulated with 40 μM PPACK and 5 U/ml clinical-grade heparin and labelled with DiOC[was drawn over a Cellix Vena8 biochip coated with fibrillar collagen (120 dynes/cm^2^, 3 min). Biochips were washed and fixed before images captured using a 40× oil immersion objective on a Leica SPE single channel confocal laser scanning microscope attached to a Leica DMi8 inverted epifluorescence microscope. Accumulation of cpdm/cpdm platelets was reduced in comparison to wt/wt platelets resulting in a reduction in thrombus volume (n = 4 mean ± s.e.). For blotting data 2-way ANOVA (variable 1 = genotype, variable 2 = agonist stimulation) followed by Bonferroni’s multiple comparison test was used to test statistical significance. For *in* vitro flow data a Mann Whitney test was used to test statistical significance. α = 0.05. Three randomly chosen fields of view were analysed per sample. Quantification was performed using Volocity® 6.1.1 Quantitation software (Perkin Elmer Inc., San Jose, CA, USA). Representative images were captured using ImageJ 1.52k (Z-projection). α = 0.05. *: p<0.05, **: p<0.01.

### Accumulation on collagen under shear is reduced in cpdm/cpdm platelets

The ability of platelets to form stable aggregates; thrombi, under shear plays a critical role in both haemostasis and thrombosis. To evaluate whether thrombus formation is altered in cpdm/cpdm mice, we tested the ability of cpdm/cpdm platelets to adhere to fibrillar collagen under arterial shear. In whole blood under non-coagulating conditions, the accumulation of cpdm/cpdm platelets was reduced in comparison to wt/wt platelets resulting in a reduction in thrombus volume (**Fig. 1C**).

### Altered haematology in cpdm/cpdm mice

It is well established that the white blood cells in the peripheral blood increase dramatically with disease progression in cpdm/cpdm mice ^21,22^, predominantly due to increases in granulocytes (eosinophils, basophils, and neutrophils). We confirm an increase in the white blood cell count (**Supp. Table 1**) in cpdm/cpdm mice without alterations in the red blood cell counts. However in contrast to a previous report^21^, platelet count was moderately decreased and mean volume increased in cpdm/cpdm mice (n = 13, p = 0.0046 and p<0.0001 respectively, Student’s t-test). These alterations in platelet volume and count were accompanied by a minor reduction in mean platelet surface expression of integrin α2 (96%), integrin β1 (86%) and GPVI (87%), however only the reduction in integrin β1 reached significance (n = 6, p = 0.0313, Wilcoxon test).

### Platelet function in cpdm/cpdm mice is reduced in response to CRP-XL

To assess whether the observed reduction in cpdm/cpdm platelet accumulation under shear reflects an underlying platelet function defect, agonist-mediated activation of integrin α_IIb_β_3_ activation and α-granule secretion was examined by flow cytometry. Activation of integrin α_IIb 3_ activation and exposure of P-selectin on the platelet surface was not significantly altered in cpdm/cpdm mice in response to a range of concentrations of the platelet activator thrombin (**Fig. 2A, B**). In contrast integrin α_IIb_β_3_ activation and P-selectin exposure induced by the GPVI agonist CRP-XL was significantly reduced in cpdm/cpdm platelets (**Fig. 2C, D**). CRP-mediated platelet function is highly dependent on secondary mediators such as ADP and TxA_2_ and therefore impaired release and/or signalling could underlie the defect in CRP-induced platelet activation. To block TxA_2_ and ADP contributions, we evaluated the effect of CRP on cpdm/cpdm platelets in the presence of the cyclooxygenase inhibitor indomethacin and the P2Y_12_ receptor antagonist ARC66096. Stimulation of platelets with CRP-XL in the presence of these inhibitors reduced both integrin α_IIb_β_3_ activation and α-granule secretion in wild-type (+/+) and cpdm/cpdm platelets (**Fig. 2C, D**). However, we observed that responses in cpdm/cpdm platelets were still significantly reduced in comparison to wild-type, indicating direct impairment of CRP-mediated platelet activation (**Fig. 2C, D**). Interestingly defective CRP-mediated integrin α_IIb_β_3_ activation and α-granule secretion in cpdm/cpdm platelets were largely restored by exogenous ADP, but not by the TxA_2_ receptor analogue U46619 (**Supp. Fig. 1**). This suggests that diminished ADP release from platelet secretory granules further contributes to the phenotype. Accordingly, we confirmed that while thrombin induced similar levels of release of the δ-granule cargo ATP in both wild-type and cpdm/cpdm platelets (**Fig. 2E**), CRP-XL-induced δ-granule release in cpdm/cpdm platelets was significantly impaired (**Fig. 2F**).

**Figure 2.**
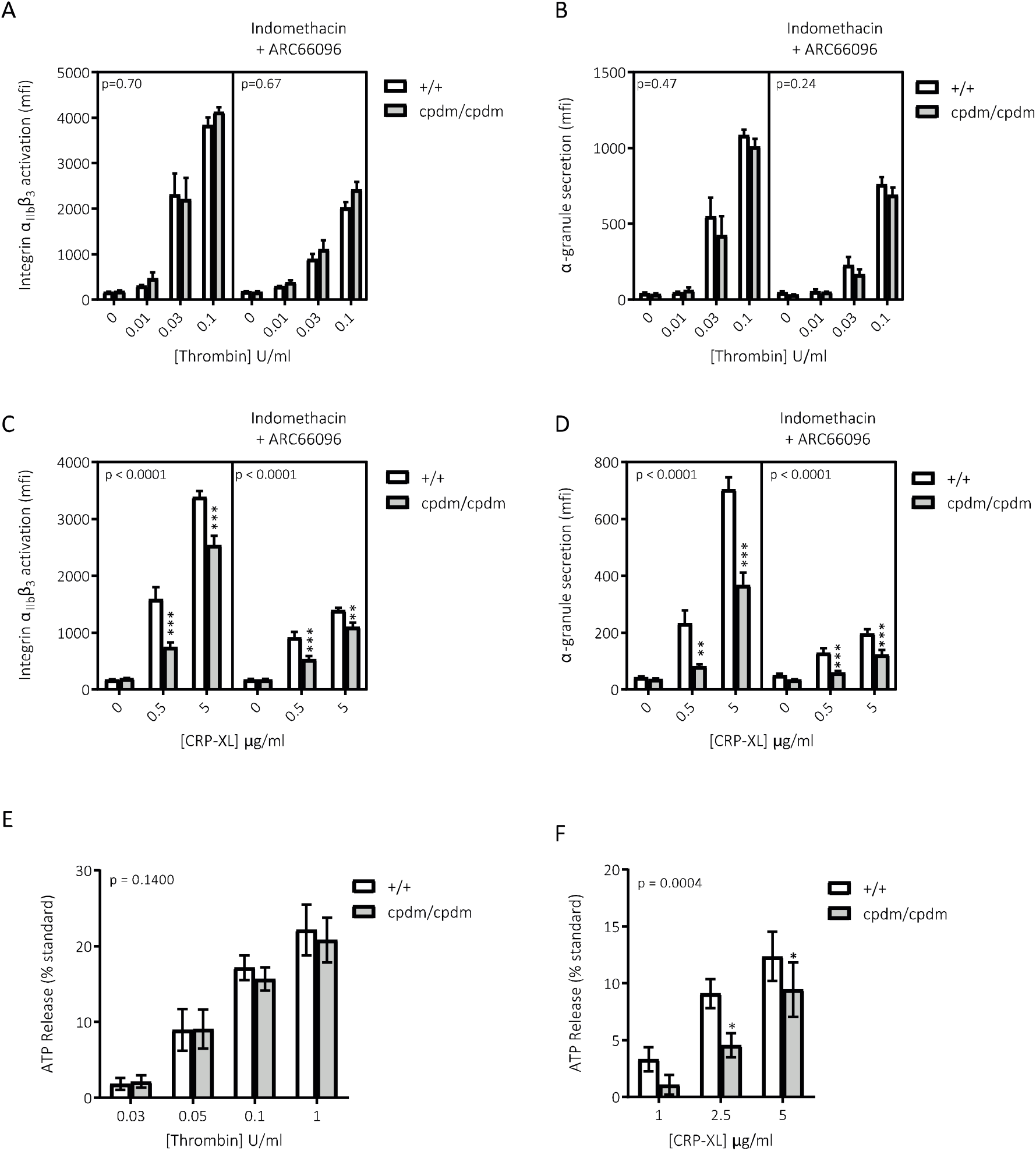
CRP-XL but not thrombin-mediated platelet activation is reduced in cpdm/cpdm platelets. **A – D)** Washed wild-type and cpdm/cpdm platelets, +/- indomethacin (10 µM) and ARC66096 (1 µM), were stimulated for 10 min with thrombin (n = 11 mean ± s.e.) or CRP-XL (n = 11 mean ± s.e.) in the presence of PE-conjugated JON/A antibody directed against the high affinity form of integrin αIIbβ3 (A, C) and FITC-conjugated (Wug.E9) antibody (B, D) against the α-granule marker CD62P (P-selectin) before fixation and analysis. **E & F)** ATP release from washed wild-type and cpdm/cpdm platelets was monitored for 5 minutes in response to indicated concentrations of thrombin (n = 6 mean ± s.e.) or CRP-XL (n = 4 mean ± s.e.). Data expressed as % of ATP standard. Results were analysed by 2-way ANOVA with Bonferroni’s post-test. P values reported for genotype variable. α = 0.05. Star notation indicates individual post-test comparisons by genotype. **: p<0.01, ***: p<0.001.

### The platelet response to U46619 is reduced in cpdm/cpdm mice

The inability of U46619 to rescue the cpdm/cpdm defect in platelet function led us to examine whether U46619-mediated platelet activation was itself compromised in cpdm/cpdm platelets. U46619 alone does not enhance integrin α_IIb_β_3_ activation and P-selectin exposure under the conditions used for FACS assays. However, a U46619-dependent component to platelet activation is clearly evident when co-incubated with 10 µM ADP, that is significantly reduced in SHARPIN-deficient platelets, despite ADP-stimulated integrin activation and P-selectin expression being unaltered (**Fig. 3A, B**). A similar trend for a reduction of both integrin α_IIb_β_3_ activation and P-selectin exposure was observed in the U46619 dose-response of cpmd/cpdm platelets co-incubated with a smaller, sub-maximal (1 µM) concentration of ADP **(Supp Fig.1C, D)**. In addition, dense granule release was impaired in U46619/ADP-stimulated platelets (**Fig 3C**). These data strongly suggest that, as with CRP, U46619-mediated platelet function is impaired in cpmd/cpdm platelets.

**Figure 3.**
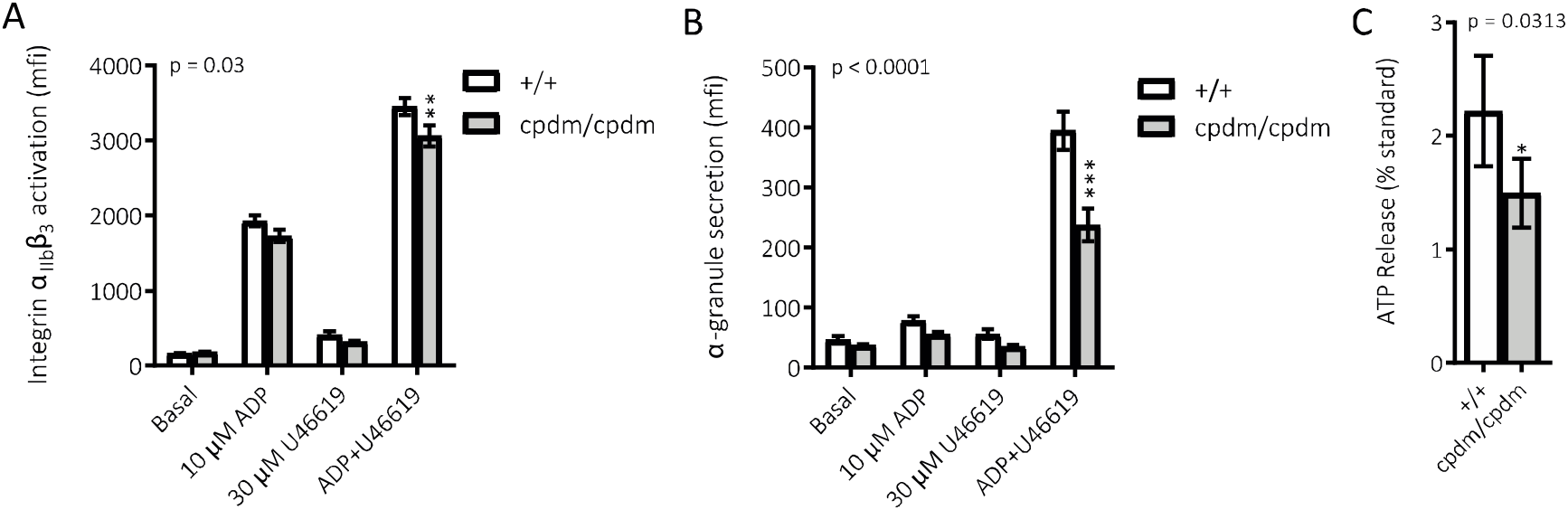
Combined U46619 and ADP give reduced activation of cpdm/cpdm platelets. Washed platelets were stimulated with the indicated agonists for 10 min in the presence of of PE-conjugated JON/A antibody directed against the high affinity form of integrin αIIbβ3 (A, C) and FITC-conjugated (Wug.E9) antibody (B, D) against the α-granule marker CD62P (P-selectin) before fixation and analysis. **A & B)** Integrin activation and α-granule secretion induced by ADP (10 µM) plus U46619 (30 µM) (n=7 mean ± s.e.). **C)** ATP release from washed wild-type and cpdm/cpdm platelets was monitored for 5 minutes in response to ADP (10µM)+U46619 (30µM) (n = 6 mean ± s.e.). Results in A - C were analysed by 2-way ANOVA with Bonferroni’s post-test. P values reported for genotype variable. α = 0.05. Star notation indicates individual post-test comparisons by genotype. Results in C were analysed by Wilcoxon test *: p<0.05, **: p<0.01, ***: p<0.001.

### GPVI and U46619-mediated signalling is altered in cpdm/cpdm platelets

A major driver of platelet δ-granule secretion is the mobilisation of intracellular calcium, therefore we examined whether CRP-XL- and U46619-mediated calcium flux was compromised in cpdm/cpdm platelets. We found a trend towards a reduction in CRP-XL-mediated calcium mobilisation (**Fig. 4A**) in cpdm/cpdm platelets, but this did not reach significance. Downstream signalling was also predominantly intact with the phosphorylation of the PI3K effector Akt and PKC substrate pleckstrin unaltered (**Fig. 4B and Supp. Fig. 2**). However, phosphorylation of MAPK ERK1/2 was significantly reduced in cpdm/cpdm platelets (**Fig. 4B**). In keeping with our findings, signalling evoked by thrombin was unaltered in cpdm/cpdm platelets (**Fig. 4B**), as was that for ADP **(Supp. Fig. 2**). Interestingly, U46619-induced calcium mobilisation was unaffected in cpdm/cpdm platelets (**Fig. 4C**), but intracellular signalling was severely impaired; phosphorylation of Akt, pleckstrin and ERK1/2 were all substantially reduced (**Fig. 4D**). Overall, these data indicate that initial calcium signalling through the TP receptor is similar in wild-type and cpdm/cpdm platelets, but more distal signalling pathways are altered, effects which may in part be due to the reduction in δ-granule secretion.

**Figure 4.**
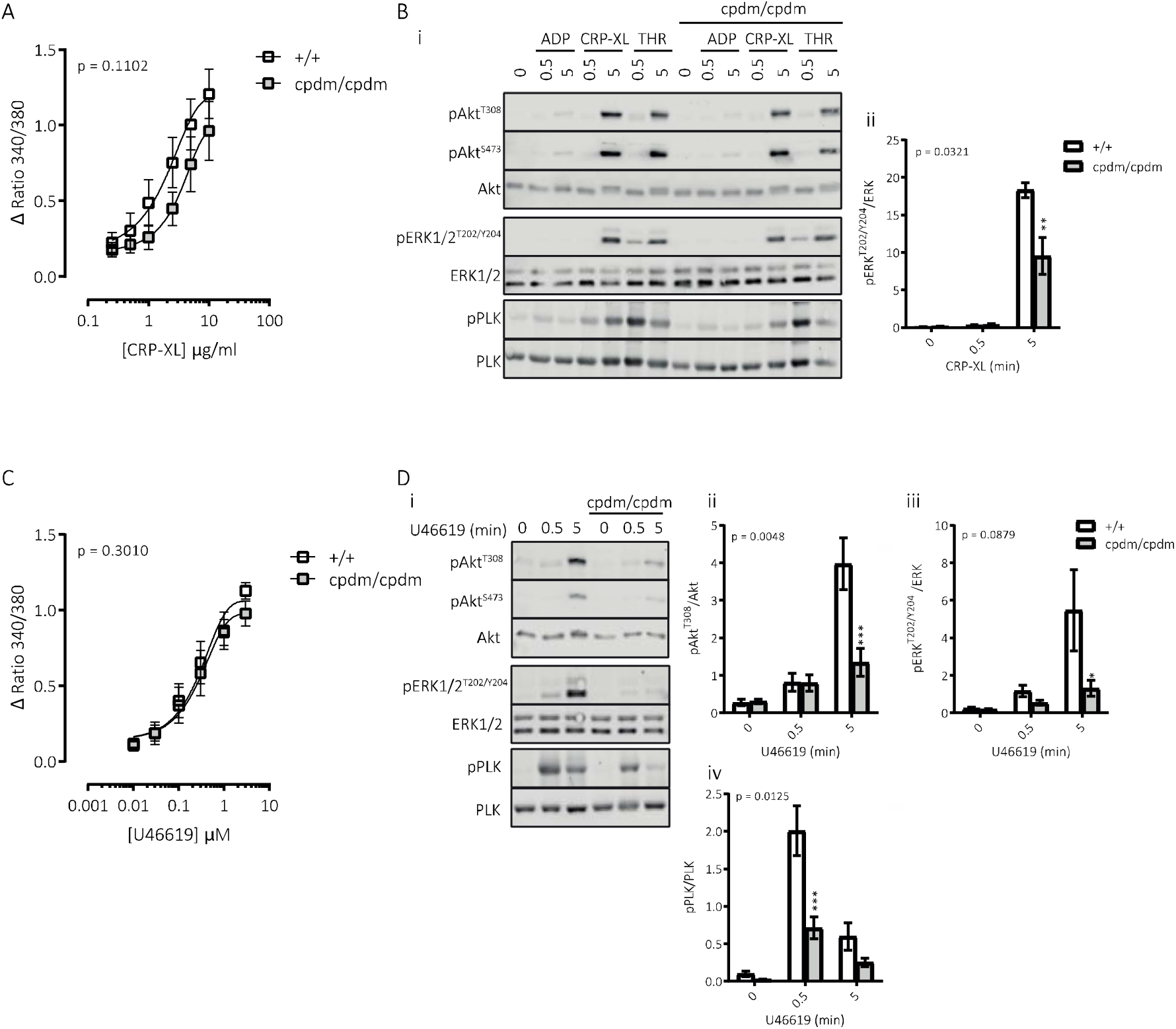
Signalling alterations in cpdm/cpdm platelets downstream of GPVI and TP receptors. **A)** Wild-type versus cpdm/cpdm Fura-PE3-loaded platelet [Ca^2+^]_i_ flux in response to indicated concentrations of CRP-XL, expressed as the change in ratio of 510 nm fluorescence emission when excited at 340 versus 380 nm (n = 4 mean ± s.e.). **B)** Western blot of washed platelets treated with ligands as indicated for 0.5 or 5 mins, before lysis and blotting for total and indicated phosphorylated species of Akt, ERK1/2 and pleckstrin. CRP-XL was used at 2 μg/ml, thrombin 0.5U/ml and ADP at 10 μM (n = 5-7, Bii mean ± s.e.). **C)** Platelet [Ca^2+^]_i_ induced by U46619 (n = 4 mean ± s.e.). **D)** Western blot of platelets stimulated with U46619 (30 μM) as indicated, followed by lysis and blotting for total and indicated phosphorylated species of Akt, ERK1/2 and pleckstrin (n = 5-7 mean ± s.e.). Results were analysed by 2-way ANOVA with Bonferroni’s post-test. P values reported for genotype variable. α = 0.05. Star notation indicates significance for individual post-test comparisons by genotype. **: p<0.01, ***: p<0.001.. *: p<0.05, **: p<0.01, ***: p<0.001.

### Activation of human platelets induces IKK-dependent phosphorylation but not degradation of IκBα

The ubiquitin ligase activity of LUBAC is required for efficient activation of canonical IKK leading to the activation of NfκB. Previous work in platelets has demonstrated the presence and activation of the IKK/NfκB pathway downstream of both inflammatory mediators (CD40L, Pam3CSK4, LPS^23,24^) and traditional platelet agonists (ADP, thrombin, collagen^10,12^) ^25,26^. A conclusive picture of IKK/NfκB regulation of platelet function has yet to form, but in general a positive role has been proposed^26^. Therefore, we examined whether inhibition of IKK/NfκB could recapitulate the phenotype observed in platelets from cpdm/cpdm mice. The major downstream substrate of the IκB kinase complex is IκBα which is phosphorylated and targeted for proteasome-mediated degradation. This degradation of IκBα is required for NfκB translocation/activation in nucleated cells. Stimulation of human platelets with either CRP-XL or thrombin caused weak phosphorylation of IκBα at Ser32/36 (**Fig. 5A and B**). However, this did not lead to a loss in total IκBα suggesting that in platelets under these conditions it is not catabolised by the proteasome. In contrast, phosphorylation of IκBα at Ser32/36 and its degradation was clearly observed in TNFα-stimulated HeLa cells. The IKK/NfκB pathway inhibitors BAY 11-7082, BMS345541, BI605906 and takinib blocked the phosphorylation of IκBα in platelets (**Fig. 5A and B**). In mouse platelets, despite significant effort, we were unable to detect phosphorylation or catabolism of IκBα by either CRP-XL or thrombin (**Supp. Fig 3**).

**Figure 5.**
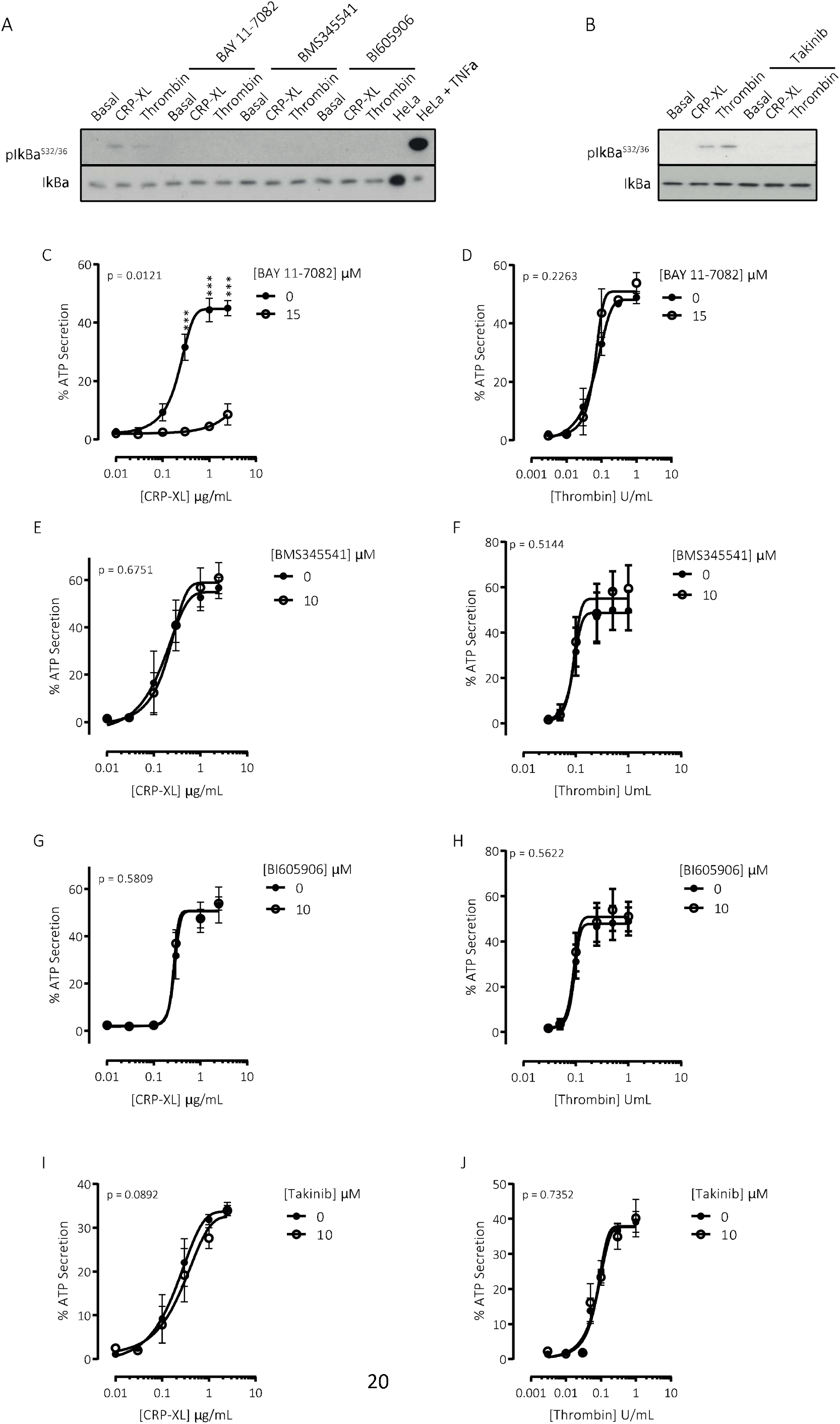
Phosphorylation of IκBα in platelets does not result in its catabolism by the proteasome and CRP-mediated δ-granule secretion in human platelets is independent of IKK activity. **A & B)** Western blot of human platelets pre-treated for 10 min with compounds indicated followed by stimulation for 5 min with (5 µg/ml) or thrombin (0.5 U/ml), before lysis and blotting for total and phosphorylated IκBα. Immunoblots representative of at least three independent experiments. **C – J)** ATP secretion from human platelets pre-treated with compounds as indicated for 10 mins followed by stimulation for 5 minutes with either 0.01 – 2 µg/ml CRP-XL (**C, E, G, I**) or 0.003 – 1 U/ml thrombin (**D, F, H, J**). BAY 11-7082; 15 µM, BMS34551; 10 µM, BI605906; 10 µM, takinib; 15 µM. Results were analysed by 2-way ANOVA (inhibitor, agonist) with Bonferroni’s post-test. P values reported for genotype variable. α = 0.05. Star notation indicates individual post-test comparisons by genotype. ***: p<0.001.

### Human platelet δ-granule secretion is dependent on ubiquitination enzyme activity but not IKK/NfκB activation

BAY 11-7082 prevents cytokine-induced IκB phosphorylation due to its inhibitory effect on the E2 conjugating enzymes; Ubc13 and UbcH7 and the E3 ligase; LUBAC^27^. As with the cpdm/cpdm mutation in mouse platelets, BAY 11-7082 treatment of human platelets reduced δ-granule secretion mediated by CRP-XL, but not thrombin (**Fig. 5C, D**). We further explored a role for IKK/NfκB using BMS345541, a highly selective IKKα/β allosteric site inhibitor^28^ previously reported to substantially inhibit granule secretion in response to low concentrations of thrombin^12^, and BI605906, a highly selective IKKβ ATP-competitive inhibitor ^29^. Crucially neither BMS345541 nor BI605906 altered δ-granule secretion in platelets induced by CRP-XL or thrombin (**Fig.5E – H**), despite blocking IκBα phosphorylation (**Fig. 5A and B**). To further confirm that δ-granule secretion is independent of IKK activity we also examined the effect of the selective inhibitor of TAK1/MAP3K7 kinase; takinib. The phosphorylation of IKKβ requires two sequential phosphorylation steps with the process initiated by TAK1 and facilitated by ubiquitination ^30^. Similar to the findings with the direct IKK inhibitors, δ-granule secretion was unaltered in the presence of takinib (**Fig. 5I, J**).

## Discussion

The results of this study demonstrate that platelets from cpdm/cpdm mice do not express the LUBAC component; SHARPIN and have reduced linear ubiquitination. Furthermore, platelets from cpdm/cpdm mice are less numerous, have a greater volume and have defects in function and signalling. However, the functional defects are not recapitulated by inhibition of the IκB kinase complex but may involve alternative signalling pathways potentially regulated by linear ubiquitination.

SHARPIN, HOIP and HOIL-1 within the LUBAC complex are key modulators of immune and inflammatory responses. Therefore, interference with how this complex can function can result in severe pathologies. This is illustrated by mouse transgenic models beyond the SHARPIN^cpdm/cpdm^ mice which have a progressive inflammatory phenotype, for example HOIP^-/-^ mice are embryonic lethal^31^ whereas HOIL-1^-/-^ mice are immunodeficient upon acute bacterial infection^32^. In extension to the mouse models, genetic alterations resulting in reduction/deletion of HOIP or HOIL-1 protein expression in humans is associated with immunodeficiency, autoinflammation and amylopectinosis^33-35^. Although no genetic alterations in SHARPIN in humans resulting in either reduction/deletion of the protein have been reported, SNPs resulting in missense variants are associated with alterations in granulocyte counts^36^. As yet platelet parameters were either not interrogated or reported in these subjects. Potentially genetic variation in LUBAC components and effectors may underlie as yet unexplained platelet function defects.

With respect to ubiquitination in platelets we detected multiple bands using an antibody specific for linear ubiquitin, furthermore the number and intensity of these bands increased upon stimulation. This is in agreement with the recent aforementioned study ^14^ where new/increased-intensity Met1 ubiquitin-positive bands were detected in NEMO immunoprecipitates from stimulated platelets. We also observed that platelets from cpdm/cpdm mice had reduced linear ubiquitination under both resting and stimulated conditions. This occurred without disruption in total ubiquitination as detected with an antibody directed against ubiquitin, polyubiquitin and ubiquitinated proteins (data not shown). Ubiquitination requires the sequential transfer of ubiquitin to a substrate by three enzymes: a ubiquitin activating enzyme (E1), a ubiquitin-conjugating enzyme (E2), and a ubiquitin ligase (E3). This can lead to either the addition of a single ubiquitin (monoubiquitination) or a chain of ubiquitin (polyubiquitination). Polyubiquitination occurs when the C-terminus of another ubiquitin links to any of the seven internal lysine residues on a previously added ubiquitin molecule. In contrast, linear (Met1-linked) ubiquitination occurs through linkage via the N-terminal methionine. At present LUBAC is the only known E3 ligase to form linear ubiquitin chains. We can therefore infer that it is the disruption of LUBAC in cpdm/cpdm platelets that directly results in a reduction in linear ubiquitination.

A major effector of linear ubiquitination in other cells is the IKK/IκBα/NfκB pathway, with linear ubiquitination ultimately leading to degradation of IκBα, freeing NfκB to translocate to the nucleus and activate transcription. Platelets are anucleate cells but do express the components of this pathway. In human platelets but not mouse we detected weak phosphorylation of IκBα in response to CRP-XL and thrombin. In both human and mouse platelets, stimulation did not lead to catabolism of IκBα. This data and the finding that inhibitors specifically targeting this pathway did not alter platelet function led us to conclude that this pathway in platelets is weakly activated and dispensable for conventional platelet function.

In contrast, pre-treatment of platelets with BAY 11-7082 an inhibitor of cytokine-induced IκB phosphorylation resulted in a reduction in CRP-XL but not thrombin-mediated platelet function. This inhibitor in contrast to the others has been demonstrated to inhibit the E2 conjugating enzymes; Ubc13 and UbcH7 and the E3 ligase; LUBAC [29]. That BAY 11-7082 pre-treatment recapitulates the effect of SHARPIN deletion therefore indicates that ubiquitination and more specifically linear (Met1-linked) ubiquitination supports platelet function in response to certain stimuli. Regarding LUBAC effectors beyond the IKK complex, as yet there are only a handful of linear ubiquitinated substrates identified. These include RIPK1^37^, RIPK2^38^, BCL10^39^ and the conjugation of linear ubiquitin chains to existing Lys63-Ub (and possibly other linkages), generating hybrid chains^40^. Linear ubiquitination of these substrates has principally been studied in the context of IKK/IκBα/NfκB signalling and therefore may not be relevant to our findings. Potentially these or other unknown linear ubiquitin substrates could be involved in regulating platelet function.

We cannot conclusively determine whether the alterations observed in platelets from cpdm/cpdm mice are driven by the deficiency in SHARPIN, by chronic alterations in the platelets driven by the inflammatory environment or through a combination of these two possibilities. However, proinflammatory conditions generally lead to enhancement, not impairment, of platelet function, by promoting immunothrombosis. A natural progression of this work would be to confirm our findings by analysing the effect of SHARPIN/LUBAC deficiency in a platelet-specific model utilising either the Pf4-Cre transgenic model or the newly published Gp1ba-Cre transgenic model, which avoids nonspecific deletion in other hematopoietic lineages as observed in the Pf4-Cre transgenic model^41^. Notwithstanding these limitations, our study demonstrates that SHARPIN deletion results in alterations in platelet parameters and function, that linear ubiquitination can be detected in platelets and that it supports platelet function. These data indicate that linear ubiquitination in platelets deserves further study and that genetic variation in LUBAC components may contribute to platelet dysfunction.

## Acknowledgements

This work was funded by the British Heart Foundation (FS/11/49/28751, PG/14/3/30565, RD/15/16/31758, PG/16/3/31833). We thank the University of Bristol’s Animal Services Unit (ASU) and Wolfson Bioimaging Facility. We also thank Emilia Peuhu (Turku Centre for Biotechnology, Finland), Elizabeth Aitken and Tony Walsh for technical assistance, Andrew Mumford, Kate Burley and Sarah Westbury for consultation regarding genetic variants and the blood donors within the Biomedical Sciences building at the University of Bristol.

## Author contributions

I.H. conceived the original idea and supervised the project. S.F.M. and I.H. conceived and planned the experiments. S.F.M., X.Z., S.M and J.L.H. performed experiments and analysed the data. S.F.M., A.W.P, S.J.M., J.L.H., and I.H. contributed to the interpretation of the results. S.F.M., J.L.H. and I.H. wrote the manuscript. All authors provided critical feedback and helped shape the research, analysis and manuscript. None of the authors has any relevant financial conflict of interest.

## References

1. Lim S, Sala C, Yoon J, et al. Sharpin, a novel postsynaptic density protein that directly interacts with the shank family of proteins. Mol Cell Neurosci. 2001;17(2):385–397.

2. Seymour RE, Hasham MG, Cox GA, et al. Spontaneous mutations in the mouse Sharpin gene result in multiorgan inflammation, immune system dysregulation and dermatitis. Genes Immun. 2007;8(5):416–421.

3. HogenEsch H, Gijbels MJ, Offerman E, van Hooft J, van Bekkum DW, Zurcher C. A spontaneous mutation characterized by chronic proliferative dermatitis in C57BL mice. Am J Pathol. 1993;143(3):972–982.

4. Tokunaga F, Nakagawa T, Nakahara M, et al. SHARPIN is a component of the NF-kappaB-activating linear ubiquitin chain assembly complex. Nature. 2011;471(7340):633–636.

5. Ikeda F, Deribe YL, Skanland SS, et al. SHARPIN forms a linear ubiquitin ligase complex regulating NF-kappaB activity and apoptosis. Nature. 2011;471(7340):637–641.

6. Dangelmaier CA, Quinter PG, Jin J, Tsygankov AY, Kunapuli SP, Daniel JL. Rapid ubiquitination of Syk following GPVI activation in platelets. Blood. 2005;105(10):3918–3924.

7. Gupta N, Li W, McIntyre TM. Deubiquitinases Modulate Platelet Proteome Ubiquitination, Aggregation, and Thrombosis. Arterioscler Thromb Vasc Biol. 2015;35(12):2657–2666.

8. Karim ZA, Vemana HP, Khasawneh FT. MALT1-ubiquitination triggers non-genomic NF-kappaB/IKK signaling upon platelet activation. PLoS One. 2015;10(3):e0119363.

9. Unsworth AJ, Bombik I, Pinto-Fernandez A, et al. Human Platelet Protein Ubiquitylation and Changes following GPVI Activation. Thromb Haemost. 2019;119(1):104–116.

10. Grundler K, Rotter R, Tilley S, et al. The proteasome regulates collagen-induced platelet aggregation via nuclear-factor-kappa-B (NFkB) activation. Thromb Res. 2016;148:15–22.

11. Spinelli SL, Casey AE, Pollock SJ, et al. Platelets and megakaryocytes contain functional nuclear factor-kappaB. Arterioscler Thromb Vasc Biol. 2010;30(3):591–598.

12. Karim ZA, Zhang J, Banerjee M, et al. IkappaB kinase phosphorylation of SNAP-23 controls platelet secretion. Blood. 2013;121(22):4567–4574.

13. Rantala JK, Pouwels J, Pellinen T, et al. SHARPIN is an endogenous inhibitor of beta1-integrin activation. Nature cell biology. 2011;13(11):1315–1324.

14. Kasirer-Friede A, Tjahjono W, Eto K, Shattil SJ. SHARPIN at the nexus of integrin, immune, and inflammatory signaling in human platelets. 2019.

15. Moore SF, Hunter RW, Harper MT, et al. Dysfunction of the PI3 kinase/Rap1/integrin alpha(IIb)beta(3) pathway underlies ex vivo platelet hypoactivity in essential thrombocythemia. Blood. 2013;121(7):1209–1219.

16. Moore SF, van den Bosch MT, Hunter RW, Sakamoto K, Poole AW, Hers I. Dual regulation of glycogen synthase kinase 3 (GSK3)alpha/beta by protein kinase C (PKC)alpha and Akt promotes thrombin-mediated integrin alphaIIbbeta3 activation and granule secretion in platelets. J Biol Chem. 2013;288(6):3918–3928.

17. Durrant TN, Hutshinson JL, Heesom KJ, et al. In-depth PtdIns(3,4,5)P3 signalosome analysis identifies DAPP1 as a negative regulator of GPVI-driven platelet function. Blood Advances. 2017;1:918–932.

18. Walsh TG, Harper MT, Poole AW. SDF-1alpha is a novel autocrine activator of platelets operating through its receptor CXCR4. Cell Signal. 2015;27(1):37–46.

19. Hunter RW, Hers I. Insulin/IGF-1 hybrid receptor expression on human platelets: consequences for the effect of insulin on platelet function. J Thromb Haemost. 2009;7(12):2123–2130.

20. Sasaki Y, Sano S, Nakahara M, et al. Defective immune responses in mice lacking LUBAC-mediated linear ubiquitination in B cells. Embo j. 2013;32(18):2463–2476.

21. Liang Y. Chronic Proliferative Dermatitis in Mice: NFkappaB Activation Autoinflammatory Disease. Patholog Res Int. 2011;2011:936794.

22. Gurung P, Sharma BR, Kanneganti TD. Distinct role of IL-1beta in instigating disease in Sharpincpdm mice. Sci Rep. 2016;6:36634.

23. Kojok K, Akoum SE, Mohsen M, Mourad W, Merhi Y. CD40L Priming of Platelets via NF-kappaB Activation is CD40- and TAK1-Dependent. J Am Heart Assoc. 2018;7(23):e03677.

24. Rivadeneyra L, Carestia A, Etulain J, et al. Regulation of platelet responses triggered by Toll-like receptor 2 and 4 ligands is another non-genomic role of nuclear factor-kappaB. Thromb Res. 2014;133(2):235–243.

25. Fuentes E, Rojas A, Palomo I. NF-kappaB signaling pathway as target for antiplatelet activity. Blood Rev. 2016;30(4):309–315.

26. Mussbacher M, Salzmann M, Brostjan C, et al. Cell Type-Specific Roles of NF-kappaB Linking Inflammation and Thrombosis. Front Immunol. 2019;10:85.

27. Strickson S, Campbell DG, Emmerich CH, et al. The anti-inflammatory drug BAY 11-7082 suppresses the MyD88-dependent signalling network by targeting the ubiquitin system. Biochem J. 2013;451(3):427–437.

28. Du Z, Whitt MA, Baumann J, et al. Inhibition of type I interferon-mediated antiviral action in human glioma cells by the IKK inhibitors BMS-345541 and TPCA-1. J Interferon Cytokine Res. 2012;32(8):368–377.

29. Clark K, Peggie M, Plater L, et al. Novel cross-talk within the IKK family controls innate immunity. Biochem J. 2011;434(1):93–104.

30. Zhang J, Clark K, Lawrence T, Peggie MW, Cohen P. An unexpected twist to the activation of IKKbeta: TAK1 primes IKKbeta for activation by autophosphorylation. Biochem J. 2014;461(3):531–537.

31. Peltzer N, Rieser E, Taraborrelli L, et al. HOIP deficiency causes embryonic lethality by aberrant TNFR1-mediated endothelial cell death. Cell Rep. 2014;9(1):153–165.

32. MacDuff DA, Reese TA, Kimmey JM, et al. Phenotypic complementation of genetic immunodeficiency by chronic herpesvirus infection. Elife. 2015;4.

33. Boisson B, Laplantine E, Dobbs K, et al. Human HOIP and LUBAC deficiency underlies autoinflammation, immunodeficiency, amylopectinosis, and lymphangiectasia. J Exp Med. 2015;212(6):939–951.

34. Boisson B, Laplantine E, Prando C, et al. Immunodeficiency, autoinflammation and amylopectinosis in humans with inherited HOIL-1 and LUBAC deficiency. Nat Immunol. 2012;13(12):1178–1186.

35. Oda H, Beck DB, Kuehn HS, et al. Second Case of HOIP Deficiency Expands Clinical Features and Defines Inflammatory Transcriptome Regulated by LUBAC. Front Immunol. 2019;10:479.

36. Astle WJ, Elding H, Jiang T, et al. The Allelic Landscape of Human Blood Cell Trait Variation and Links to Common Complex Disease. Cell. 2016;167(5):1415-1429.e1419.

37. Keusekotten K, Elliott PR, Glockner L, et al. OTULIN antagonizes LUBAC signaling by specifically hydrolyzing Met1-linked polyubiquitin. Cell. 2013;153(6):1312–1326.

38. Fiil BK, Damgaard RB, Wagner SA, et al. OTULIN restricts Met1-linked ubiquitination to control innate immune signaling. Mol Cell. 2013;50(6):818–830.

39. Satpathy S, Wagner SA, Beli P, et al. Systems-wide analysis of BCR signalosomes and downstream phosphorylation and ubiquitylation. Mol Syst Biol. 2015;11(6):810.

40. Emmerich CH, Ordureau A, Strickson S, et al. Activation of the canonical IKK complex by K63/M1-linked hybrid ubiquitin chains. Proc Natl Acad Sci U S A. 2013;110(38):15247–15252.

41. Nagy Z, Vogtle T, Geer MJ, et al. The Gp1ba-Cre transgenic mouse: a new model to delineate platelet and leukocyte functions. Blood. 2019;133(4):331–343.

