## Supplementary Figures for "A role for SHARPIN in platelet linear protein ubiquitination and function"

**Supplementary Figure 1. ADP but not U46619 can rescue the GPVI functional defect in cpdm/cpdm platelets** Washed platelets were co-stimulated with either vehicle or CRP-XL (0, 0.5, 5 μg/ml) together with either vehicle, ADP (10 μM) or U46 (30 μM) for 10 minutes in the presence of PE-conjugated JON/A antibody directed against the high affinity form of integrin αIIbβ3 **(A)** and FITC-conjugated (Wug.E9) antibody **(B)** against the α-granule marker CD62P (P-selectin) before fixation and flow cytometry n= 3, mean±s.e. Data were analysed by 2-way ANOVA (variable 1 = genotype, variable 2 = stimulation) followed by Bonferroni post-testing. α = 0.05. P values shown are those for the effect of genotype. Star notation indicates statistical significance for individual comparisons. *: p<0.05 **: p<0.01, ***: p<0.001. **C & D)** Integrin activation and α-granule secretion in response to varying concentrations of U46619 (10 nM to 10 μM) in the presence of ADP (1 μM) (n=2 mean ± s.e.).

**Supplementary Figure 2. Quantification of CRP-XL, Thrombin and ADP induced signalling in cpdm/cpdm platelets.** Washed platelets were treated with **A)** CRP-XL (2 μg/ml), **B)** thrombin (0.5U/ml) or **C)** ADP (10 μM) and lysed in 4× NuPAGE sample buffer containing 0.5 M DTT before analysis by SDS-PAGE/Western blotting as indicated (n=4-7, mean±s.e). A 2-way ANOVA (variable 1 = genotype, variable 2 = stimulation) followed by Bonferroni's multiple comparison test was used to test statistical significance. α = 0.05. The P values in the figure indicate whether a difference exists between genotypes.

**Supplementary Figure 3. Neither CRP-XL nor thrombin induce phosphorylation or degradation of IκBα in mouse platelets**. Western blot of washed wild-type and cpdm/cpdm mouse platelet lysates following treatment with ligands indicated for 5 min. HeLa lysates were used as a positive control. Immunoblot representative of at least three independent experiments.

S.Figure 1

A

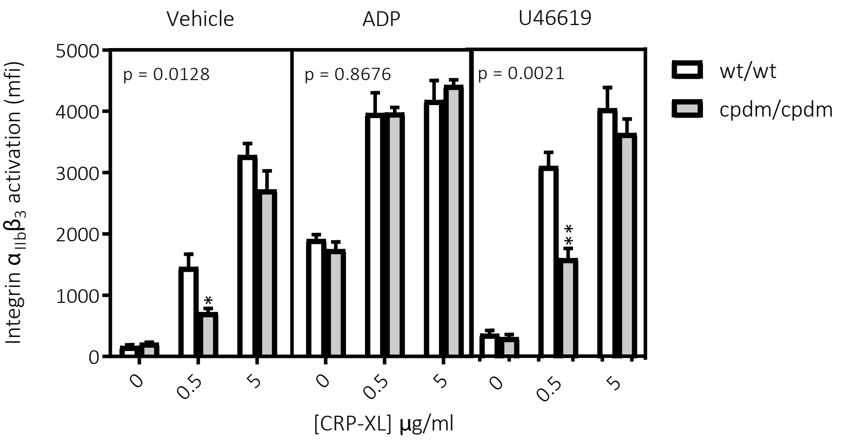


B

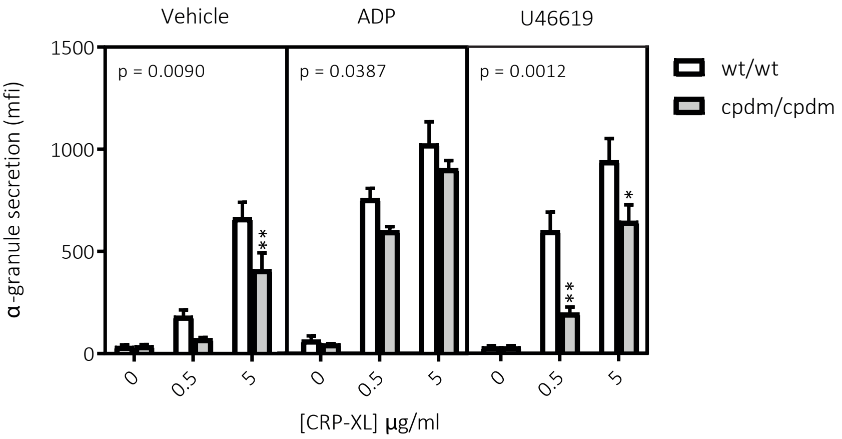


C

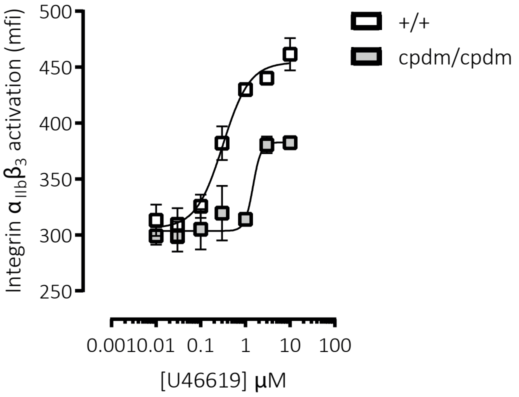


D

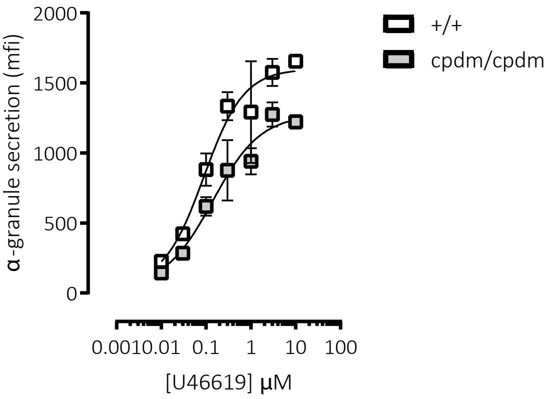

### S.Figure 2

A

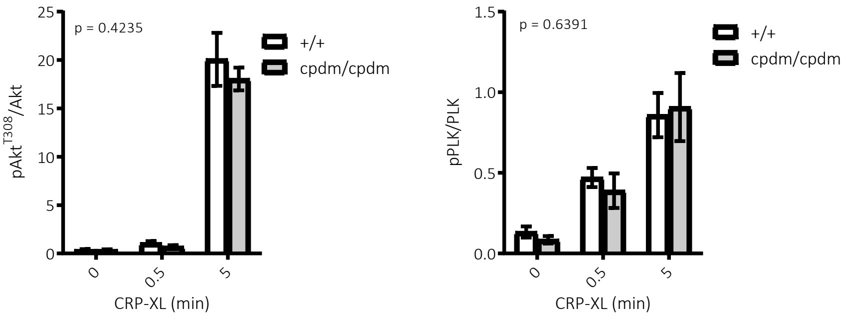


B

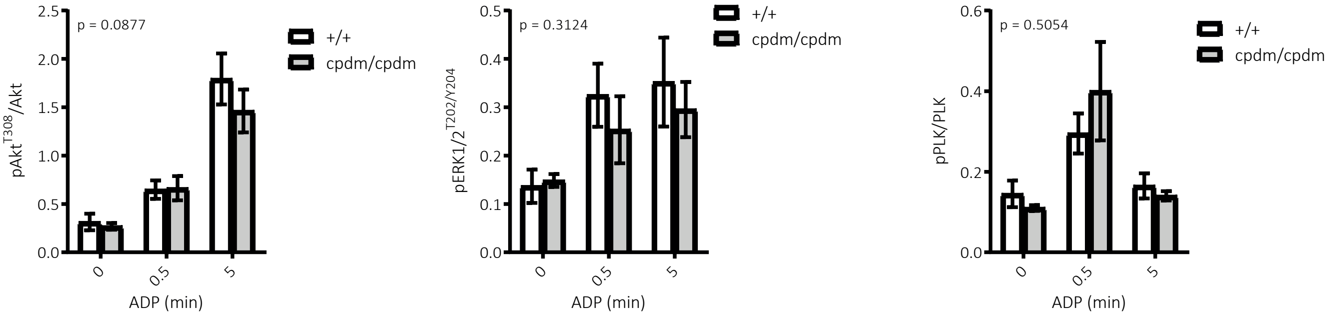


C

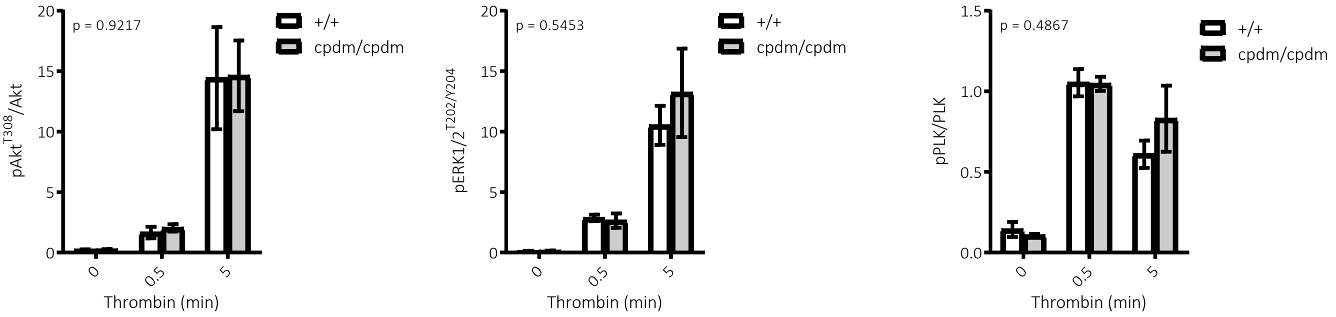

S.Figure 3

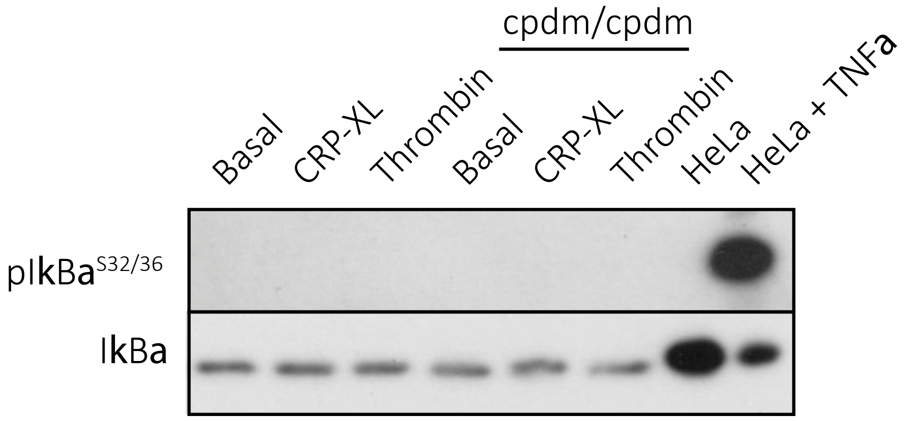
cpdm/cpdm
Basal
CRP-XL
Thrombin
Basal
CRP-XL
Thrombin
HeLa
HeLa + TNFa
pIkBa S32/36
IkBa

**S.Table 1: Whole blood counts and platelet surface receptor expression in wild-type (+/+) and Sharpin-deficient (cpdm/cpdm) mice.**

| | +/+ | | cpdm/cpdm | | | |
|---|---|---|---|---|---|---|
| | Mean | s.e.m | Mean | s.e.m | n | p-value |
| Platelets $10^3/mm^3$ | 1054 | 40 | 862 | 40 | 13 | 0.0046 |
| MPV $mm^3$ | 5.23 | 0.05 | 5.78 | 0.04 | 13 | <0.0001 |
| WBC $10^3/mm^3$ | 11.0 | 0.6 | 58.7 | 3.2 | 13 | <0.0001 |
| RBC $10^6/mm^3$ | 11.1 | 0.2 | 10.6 | 0.2 | 13 | 0.024 |
| Integrin $\alpha_{II}$ mfi | 4597 | 464 | 4448 | 504 | 6 | 0.5625 |
| Integrin $\beta_3$ mfi | 4948 | 142 | 5377 | 427 | 6 | 0.4375 |
| Integrin $\alpha_2$ mfi | 582 | 20 | 560 | 28 | 6 | 0.0938 |
| Integrin $\beta_1$ mfi | 2650 | 125 | 2289 | 66 | 6 | 0.0313 |
| GPVI mfi | 966 | 8.9 | 843 | 14.4 | 5 | 0.0625 |

Complete blood counts were conducted using a Pentra ES60 (Horiba)and adjusted for anticoagulant volume. Surface expression of integrin $\alpha_{IIb}$ (CD41), integrin $\beta_3$ (CD61), integrin $\alpha_2$ (CD49b), integrin $\beta_1$ (CD29) and GPVI were measured in resting platelets by flow cytometry using FITC-conjugated antibodies. Student's t-tests (haematology) and Wilcoxon tests (surface glycoprotein) were used to test for statistical differences between genotypes.
